# The role of histone modifications and transposable elements in the epigenetic regulation of gene dosage after gene duplication

**DOI:** 10.1101/2024.11.18.624187

**Authors:** Thomas D Wolfe, Frederic J.J. Chain

## Abstract

The duplication of genes has long been recognized as a substrate for evolutionary novelty and adaptation, but the factors that govern fixation of paralogs soon after duplication are only partially understood. Duplication often leads to an increase in gene dosage, or the amount of functional gene product. For genes with which an increased dosage is harmful (i.e., triplosensitive genes), a dosage balancing mechanism needs to be present immediately after duplication if it is to evade negative selection. Previous research in vertebrates has demonstrated a potential role for epigenetic factors in allowing triplosensitive genes to increase in copy number by regulating their expression post-duplication. Here we expand this research by investigating the epigenetic landscape of duplicate genes in *D. discoideum*, a basal lineage separated from humans by over a billion years. We found that activating histone modifications are quickly lost in duplicate genes before gradually increasing in enrichment as paralogs age. For the repressive modification H3K9me3, we found it was enriched in the youngest paralogs, and that this enrichment was likely mediated by heterochromatin spread from transposable elements. We similarly found enrichment of H3K9me3 in young human duplicates, and again found transposable elements as a potential mediator. Finally, we leveraged recent genome-wide estimates of triplosensitivity in human genes to directly examine the relationship between this kind of dosage sensitivity and enrichment for repressive histone modifications. Interestingly, while we found no significant link between enrichment for the repressive mark H3K9me3 and triplosensitivity in human paralogs, we did find a significant association between triplosensitivity and transposon proximity. Our findings suggest that transposons may contribute to the epigenetic regulatory environment associated with dosage balancing of young duplicates in both protists and humans.

## Introduction

Gene duplication is an important mechanism in generating the raw material for the emergence of evolutionary novelty and adaptation at the molecular level across all branches of life (Katju and Bergthorsson 2013; Lynch and Conery 2000). From conserved processes like homologous recombination in DNA repair, which requires the use of several paralogs of the conserved Rad51 protein in both yeast and humans to function properly (Roy et al. 2021; Greenhough et al. 2023), to critical systems in vertebrates like oxygen transport and storage (Storz 2016), gene duplications have provided the substrate for evolution to act on. There is also evidence to suggest that gene duplications have been critical in the development of human-specific higher-order cognitive faculties (Zimmer and Montgomery 2015; Sassa 2013). Yet, while the mechanisms behind the initial duplication of genes are fairly well understood, it remains unclear to what extent different evolutionary models explain the subsequent retention and evolution of gene duplicates (Innan and Kondrashov 2010; Kuzmin et al. 2022). One challenge in formulating a working model of paralog evolution is accounting for the survival of newly formed duplicates despite the potential for a harmful increase in gene dosage. While paralogs born of a whole-genome duplication are less likely to exhibit maladaptive dosage effects, as each gene is duplicated along with all interacting partners, smaller scale duplications may result in the duplication of only one member of a protein complex or dosage-sensitive network thus leading to a stoichiometric imbalance.

Additionally, some genes are prone to promiscuous binding, making their duplication potentially harmful to the functioning of other proteins (Vavouri et al. 2009). While positive selection may act on some new duplicate genes for which an increased dosage is beneficial (Kondrashov 2012), it has been shown in multiple yeast and mammalian species that lineage-specific paralogs typically have an average expression lower than that of an orthologous singleton, suggesting that combined gene dosage among young paralogs is reduced to a level closer to that of the ancestral singleton shortly after duplication (Qian et al. 2010; Lan and Pritchard 2016). Though this may be the result of regulatory mutations acquired via drift, the fact that this expression reduction is more pronounced in dosage-sensitive genes suggests that it is a response to deleterious gene dosage increase (Qian et al. 2010; Chang and Liao 2017).

One potential mechanism underlying this dosage balancing model soon after duplication is epigenetic regulation. The recruitment or pre-existence of epigenetic modifications at duplicated loci offers an elegant explanation for the reduced expression of duplicates shortly after their emergence; epigenetic marks could be established or adjusted quickly to effect a change in expression long before the acquisition of mutations that could similarly balance gene dosage, which may take generations to accrue. To this end, reduction in expression shortly after duplication has been linked to epigenetic modifications in several animal species, wherein a relationship exists between the initial expression decrease in young duplicates and enrichment for repressive DNA methylation and histone modifications (Keller and Yi 2014; Chang and Liao 2017; Huang and Chain 2021). It is currently unclear to what extent these relationships hold in other divergent eukaryotic lineages, and what mechanism may be available to the genome that might account for this early repression of gene duplicates.

To further examine the potential role of epigenetic regulation in driving dosage balancing of duplicate genes, we used a diverse set of publicly available epigenomic and transcriptomic data from the social amoeba *Dictyostelium discoideum*. As a basal eukaryote which often serves as a model for the transition of eukaryotes from unicellular to multicellular organisms (Bozzaro 2013), *D. discoideum* is a well-studied member of the monophyletic Amoebozoa supergroup, the protist sister taxa to the proposed Obazoa clade containing all animals and fungi (Cavalier-Smith et al. 2015; Brown et al. 2013; Burki et al. 2020). While DNA methylation is present only at extremely low levels within the *D. discoideum* genome (Drewell et al. 2023), conserved histone modifications such as histone acetylation and histone methylation are present. To identify patterns of transcriptional repression and activation, we used publicly available ChIP-seq data corresponding to tri-methylation of the 4^th^ lysine on histone H3 (H3K4me3), trimethylation of the 9^th^ lysine on histone H3 (H3K9me3), acetylation of the 27^th^ lysine on histone H3 (H3K27ac), as well as ATAC-seq and RNA-seq data. We chose these modifications because H3K4me3 and H3K27ac are well established markers of active chromatin regions (Black et al. 2012; Zhao and Garcia 2015), H3K9me3 is the canonical marker of heterochromatin (Hyun et al. 2017), and ATAC-seq serves as a sensitive assay for identifying regions of accessible chromatin genome-wide (Buenrostro et al. 2015). After identifying enrichment of H3K9me3 in young duplicates proximal to transposable element (TE) sequences in *D. discoideum*, we tested for similar enrichment in humans, a species separated from *D. discoideum* by over a billion years.

## Results

### Distribution and regulatory effects of H3K9me3, H3K4me3, H3K27ac and open chromatin in *D. discoideum*

Before investigating chromatin dynamics that might indicate dosage balancing in duplicate genes, we established the distribution and association with gene expression of the histone modifications H3K9me3, H3K4me3, H3K27ac, and open chromatin regions in *D. discoideum*. Using published ChIP-seq and ATAC-seq data sampled from multiple life-cycle stages (Wang et al. 2021), we started by identifying peak areas of enrichment for these chromatin features genome-wide (see methods: ChIP-seq and ATAC-seq alignment and peak calling). We then explored the relationship between enrichment of these chromatin features and gene expression in all protein coding genes. In many organisms H3K9me3 deposition is targeted at non-coding and repetitive regions, including transposable elements (TEs) (Le Thomas et al. 2013; Pezic et al. 2014; Frapporti et al. 2019). In *D. discoideum* we found as well that H3K9me3 was strongly associated with TE sequences, with TE sequences making up 60% of all bases overlapping an H3K9me3 peak in at least one sample despite comprising < 5% of the genome (supplementary figure 1). Open reading frames of TEs correspondingly had very low expression compared to other genes (median TPM of 0.19 for TEs and 4.77 for other coding genes, supplementary figure 2). We therefore excluded the open reading frames of TEs from the group of coding genes used for this analysis, as they are likely under a different regulatory program than strictly endogenous genes.

Briefly, for each chromatin feature we calculated the frequency of peaks overlapping each base from 500 bp upstream of the transcription start site (TSS) to 500 bp downstream of the transcription termination site (TTS) as the fold change over the per-base average in all coding genes (figure 1, see methods: *ChIP-seq and ATAC-seq enrichment in D. discoideum*). After binning all genes into expression deciles, we found that enrichment for H3K4me3, H3K27ac, and ATAC-seq peaks within the gene body increased substantially as expression increased, while depletion of each peak type immediately upstream of the TSS and downstream of the TTS was consistently observed at each level of expression (figure 1a-c). The repressive mark H3K9me3 was found to be enriched in the lowest expression decile (figure 1d) with a uniform distribution across the region observed. As TEs appear to be the primary target of H3K9me3 deposition (supplementary figure 1), we then investigated the possibility that other genes with overlapping H3K9me3 peaks in at least one sample were proximal to TEs. As expected, coding genes that overlapped with an H3K9me3 peak were distributed significantly more closely to TEs than those with no H3K9me3 overlap (supplementary figure 3, *p* value < 0.001, Mann-Whitney U).

**Figure 1:**
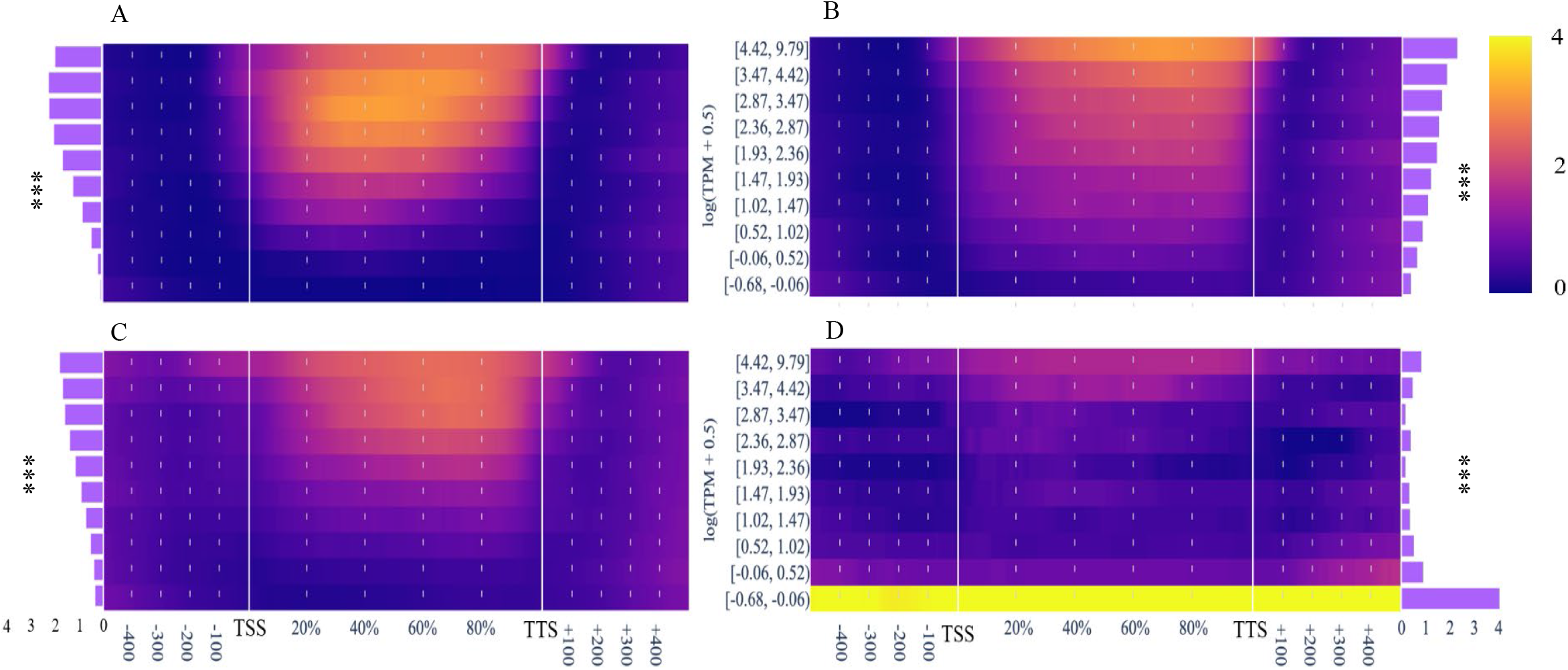
Heatmaps showing the peak fold change over the per-base average for all coding genes for four different chromatin features: (A) H3k4me3, (B) H3K27ac, (C) open chromatin (measured via ATAC-seq), and (D) H3K9me3. For each peak type, all coding genes were separated into deciles based on average expression (log-transformed with a pseudo-count of 0.5), increasing along the y-axis. Enrichment was measured from the TSS -500 bp to the TTS +500 bp (x-axis), with enrichment across the gene body interpolated into percentile intervals for each gene to create a common domain. Bar graphs on the left or right of each panel represent the fold change over the per-base average for coding genes within the predictive region of the respective chromatin feature. *** indicates a *p* value less than 0.001.

To more precisely define the relationships between enrichment for chromatin features and expression, we calculated the correlation between gene expression and peak presence at each base within the gene region specified above for each chromatin feature. For each feature we identified a contiguous span of significant correlations (after correction for multiple testing) around the TSS, and considered these regions to be most predictive of expression (table 1, see methods: ChIP-seq and ATAC-seq enrichment in *D. discoideum*). Indeed, after correction for multiple testing (Bonferroni) H3K4me3, H3K27ac, and open chromatin all increased significantly within the predictive regions (table 1, supplementary figure 4) as expression increased while H3K9me3 decreased rapidly (*p* values < 0.001, method Kruskal-Wallis, figure 1).

**Table 1:** Contiguous regions surrounding the TSS of coding genes found to be significantly correlated with expression when overlapped by a peak from one of the four chromatin features assayed. TSS: Transcription Start Site, TTS: Transcription termination site, +/-N bp: N bases upstream (-) or downstream (+) of the gene feature. Percentages were used for regions that terminated within the gene body, as the chromatin feature peaks in gene body regions of all coding genes were interpolated to percentiles to provide a common domain.

| Chromatin Feature | Region Start | Region End |
| --- | --- | --- |
| H3K27ac | TSS – 141 bp | TTS + 148 bp |
| H3K4me3 | TSS – 200 bp | TTS +164 bp |
| H3K9me3 | TSS – 500 bp | TSS + 15% |
| Open Chromatin (ATAC-seq peak) | TSS – 500 bp | TTS + 220 bp |

### Changes to chromatin in duplicate genes over evolutionary time indicate gradual activation

To populate a set of independent duplicate gene pairs, we selected pairs that shared a reciprocal highest pairwise sequence identity among the set of all likely paralogs identified by the Ensembl Compara pipeline (Howe et al. 2021) (see methods: Identifying duplicate genes and transposable elements in *D. discoideum*). To determine how the distribution of open chromatin and each histone modification changed over time post-duplication, we compared peak enrichment around predictive regions across different levels of duplicate gene divergence, as measured by the pairwise rate of synonymous mutation (dS). As with enrichment relative to gene expression, we chose the same broad region spanning 500 bases upstream of the TSS to 500 bases downstream of the TTS, and again calculated the fold-change relative to the average frequency for all bases in the region for each coding gene.

We observed an initial modest enrichment in H3K4me3, H3K27ac and ATAC-seq peaks in the lowest *dS* decile within the predictive regions, followed by a rapid decline in enrichment in the next decile, and a subsequent gradual increase in enrichment as dS increased (figure 2a-c). After correction for multiple testing (Bonferroni) the differences in enrichment across dS deciles were significant for H3K4me3 (*p* value < 0.001, Kruskal–Wallis), though they were not for any of the other chromatin features (table 2). This was particularly surprising for H3K9me3, as a clear enrichment for the mark could be seen in the two lowest *dS* deciles (figure 2d). The non-significant *p* values may have been due to the Kruskal-Wallis test being underpowered in the presence of zero-inflated data and relatively small sample sizes (∼50 per decile). Given this, we decided to compare the proportion of genes with an H3K9me3 peak overlapping the predictive region in the two lowest deciles to (a) the proportion in older duplicates, and (b) the proportion in singletons. In the two lowest deciles, we found an H3K9me3 peak overlapping 25 paralogs and 181 without an H3K9me3 overlap, a ratio significantly higher than we found among singletons or the remaining duplicate deciles (singleton ratio: 170:4805, remaining decile ratio: 28:794, *p* values both < 0.001, Fisher’s exact test).

**Figure 2:**
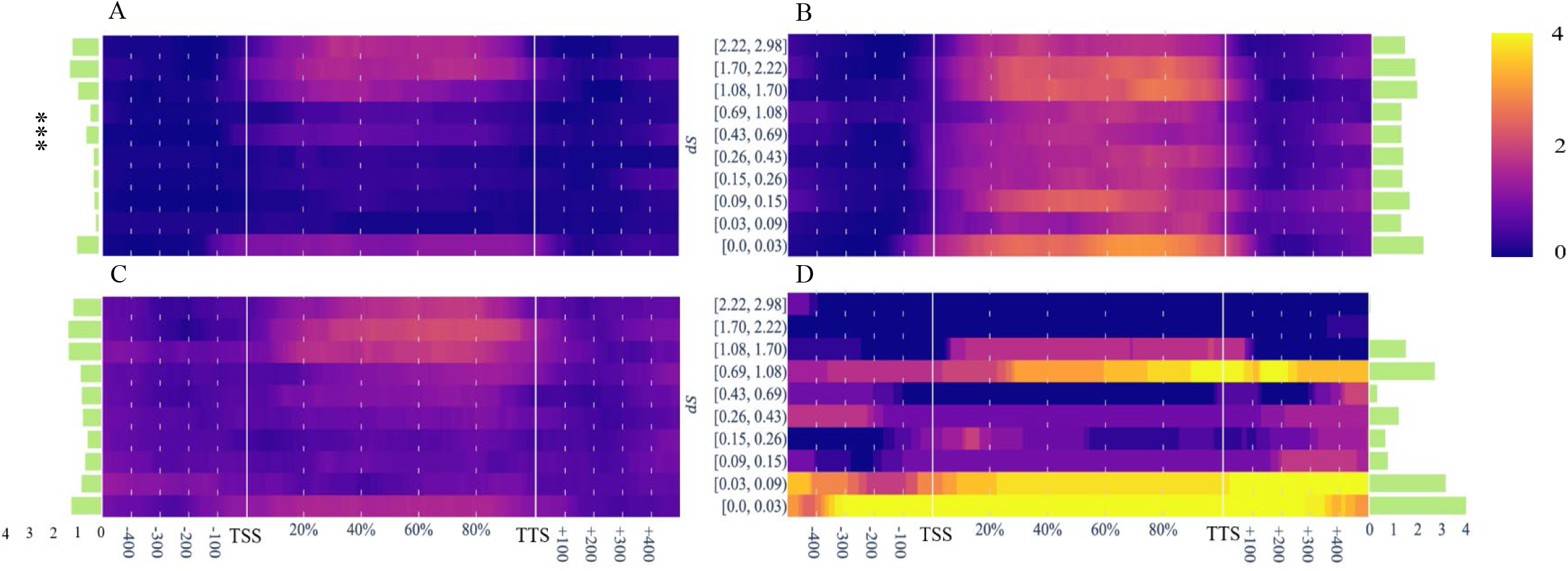
Heatmaps showing the frequency of enrichment peaks in paralogs as multiples of the per-base average for all coding genes for four different chromatin features: (A) H3K4me3, (B) H3K27ac, (C) open chromatin (measured via ATAC-seq), and (D) H3K9me3. For each peak type, all individual paralogs were separated into deciles based on the pairwise *dS* with their paralog pair, increasing along the y-axis. Enrichment was measured from 500 bases upstream of TSS to 500 baes downstream of the TTS (x-axis), with enrichment across the gene body interpolated into percentile intervals for each gene to create a common domain. Bar graphs on the right of each panel represent the frequency multiple over the per-base average for coding genes within the predictive region of the respective chromatin feature. *** indicates a *p* value less than 0.001.

**Table 2:** Results of the Kruskal-Wallis test of chromatin feature enrichment in predictive regions with respect to gene expression or rate of synonymous divergence (*dS*) between paralog pairs. The test was used for each chromatin feature with either expression or *dS* deciles used as independent samples. The Bonferroni correction was applied to the *p* values grouped into enrichment/expression tests and the enrichment/*dS* tests.

| Chromatin Feature | *p* value (Enrichment vs. Expression) | *p* value (Enrichment vs. *dS*) |
| --- | --- | --- |
| H3K27ac | ~0 | 0.33 |
| H3K4me3 | ~0 | <<0.05 |
| H3K9me3 | << 0.05 | 0.19 |
| Open Chromatin (ATAC-seq peak) | ~0 | 0.16 |

Consistent with the enrichment patterns among activating modifications, we observed an initially high level of average expression among paralog pairs, which rapidly decreased before gradually increasing to a level close to the average for all coding genes (figure 3a). Additionally, the difference in expression between each duplicate gene pair, which we calculated as the Euclidean distance of each pair member’s RNA-seq expression values across samples, increased over evolutionary time (figure 3b). It should be noted however that given the high sequency identity among the least diverged duplicates, the difference in expression may be underestimated due to ambiguous RNA-seq read mapping.

**Figure 3:**
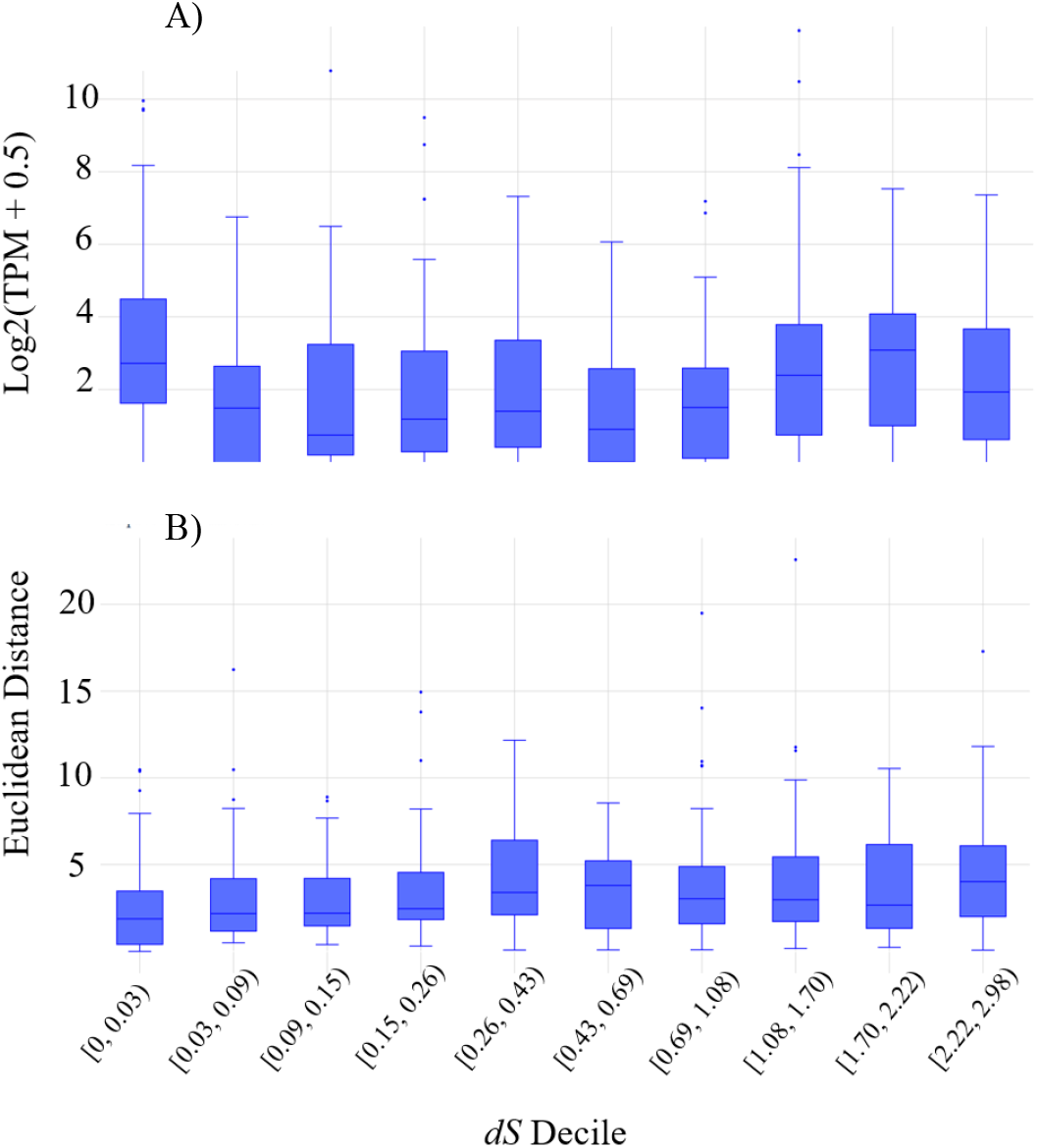
After an initially high level of expression, average paralog pair expression gradually increases. Expression divergence between airs steadily increases as paralogs diverge. (A) Distribution of average gene expression—measured as the log2-transformed transcripts per million (TPM) with a pseudocount of 0.5 added—within paralog pairs (y-axis) and (B) pairwise difference in expression (pairwise Euclidean distance of expression across life cycle stages, y-axis) for each paralog pair separated into *dS* deciles (increasing along the x-axis).

Given the repressive mark H3K9me3 was enriched in the lowest dS decile despite the high expression level, and given activating modifications were only modestly enriched in the same gene cohort, we investigated the possibility that the expression levels of some duplicates were artificially inflated. Specifically, we investigated the possibility that the more recently duplicated genes existed in a higher copy number than assembled in the reference genome, possibly leading to inflated RNA-seq counts if reads from multiple cryptic gene copies were mapping to the same locus in the reference. To account for this possibility we used the pileup of input DNA control reads from the ChIP-seq assay to adjust the RNA-seq data, as input read enrichment has been previously demonstrated to correlate well with gene copy number (Vega et al. 2009). We did this by scaling down the expression level of each gene by the ratio of reads mapped to the respective gene’s locus over the average for single-copy genes (see methods: RNA-seq alignment and transcript quantification). Even after this adjustment, a high expression level remained among the duplicates in the lowest dS decile (supplementary figure 5), with the median expression level of that cohort the highest among all deciles. Assuming that the duplicates in this group are truly highly expressed, we investigated the possibility of there being two sub-groups among the most recently duplicated genes: one with high expression and moderate enrichment for activating modifications, and one with low expression and enrichment for H3K9me3.

Indeed, the expression level of duplicates in the two lowest dS deciles that were overlapping an H3K9me3 peak was significantly lower than that of those which did not overlap a peak (*p* value < 0.001, Mann–Whitney U). Interestingly, however, most of the duplicates (18 of 25) overlapping an H3K9me3 peak were also overlapping one of the activating features, primarily ATAC-seq or H3K27ac peaks (13 of 25 and 10 of 25, respectively, figure 4). As with all coding genes in *D. discoideum*, we found that individual duplicates with H3K9me3 peaks were also inordinately close to TEs (supplementary figure 6, *p* value < 0.001, Mann–Whitney U).

**Figure 4:**
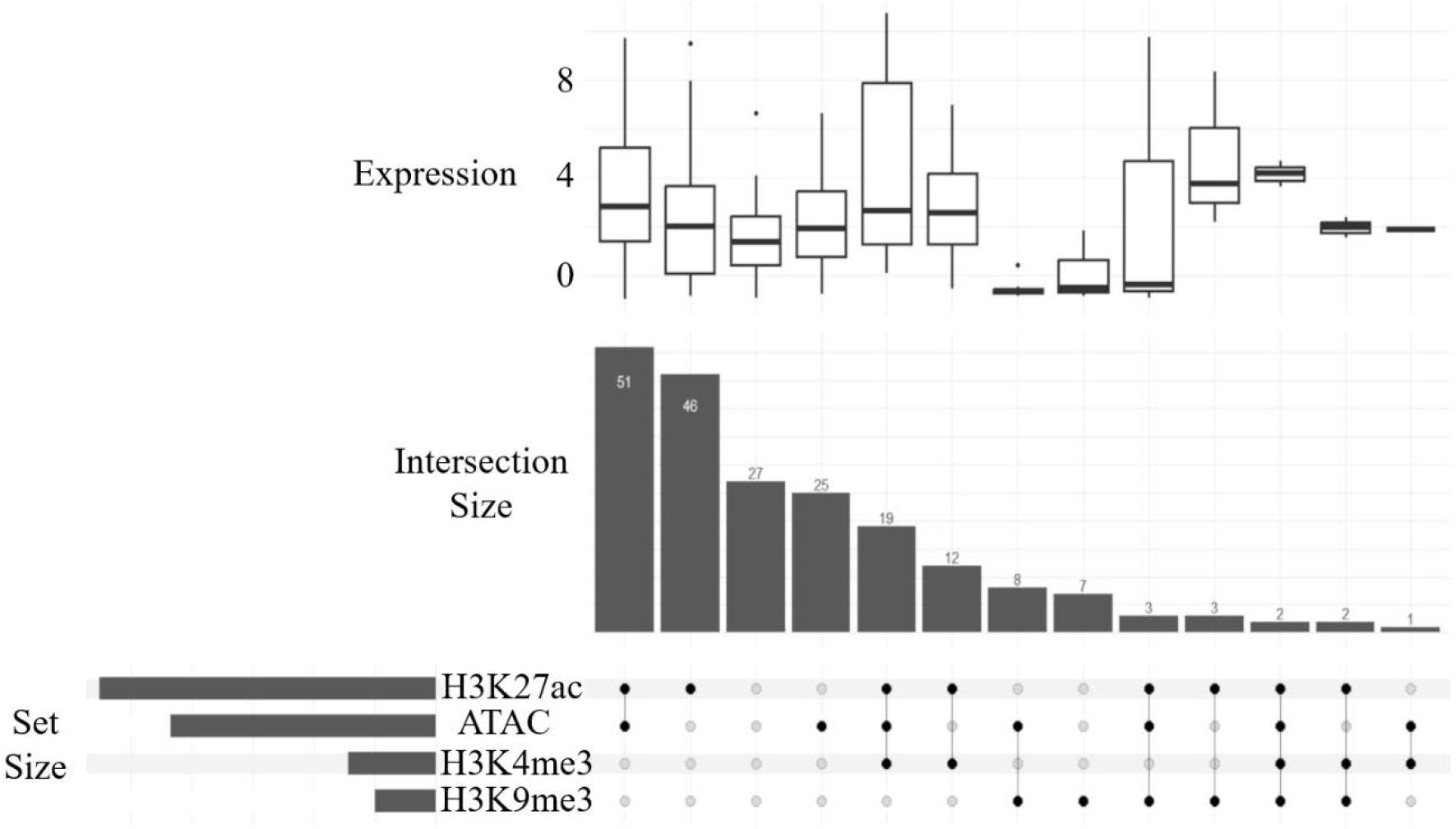
Separation of all paralog pairs in the two lowest dS deciles (0 <= dS < 0.9) into intersecting sets based on the presence of chromatin feature peaks. Box plots showing the distribution of paralog expression are placed above the respective intersection. Expression was measured as the log2-transformed TPM with a pseudocount of 0.5 added and averaged across samples for each paralog.

### H3K9me3 is also enriched in young duplicates in humans

To test if the pattern of enrichment for H3K9me3 in young duplicates is also found in animals, we used publicly available ChIP-seq data to calculate enrichment for the repressive mark in human duplicate genes. As with *D. discoideum*, we identified a set of independent duplicate pairs and calculated the fold change in H3K9me3 coverage over the average in singleton coding genes from 500 bases upstream of the TSS to 500 bases downstream of the TTS (figure 5, see methods: Identifying duplicate genes and transposable elements in humans, H3K9me3 enrichment in humans). We found that enrichment for H3K9me3 was greatest among the younger duplicate genes (0 <= dS < 0.19, figure 5), albeit highest in dS bins > 0, and that the repressive mark was enriched in humans in duplicates compared to singletons (*p* value < 0.001, Fisher’s exact test).

**Figure 5:**
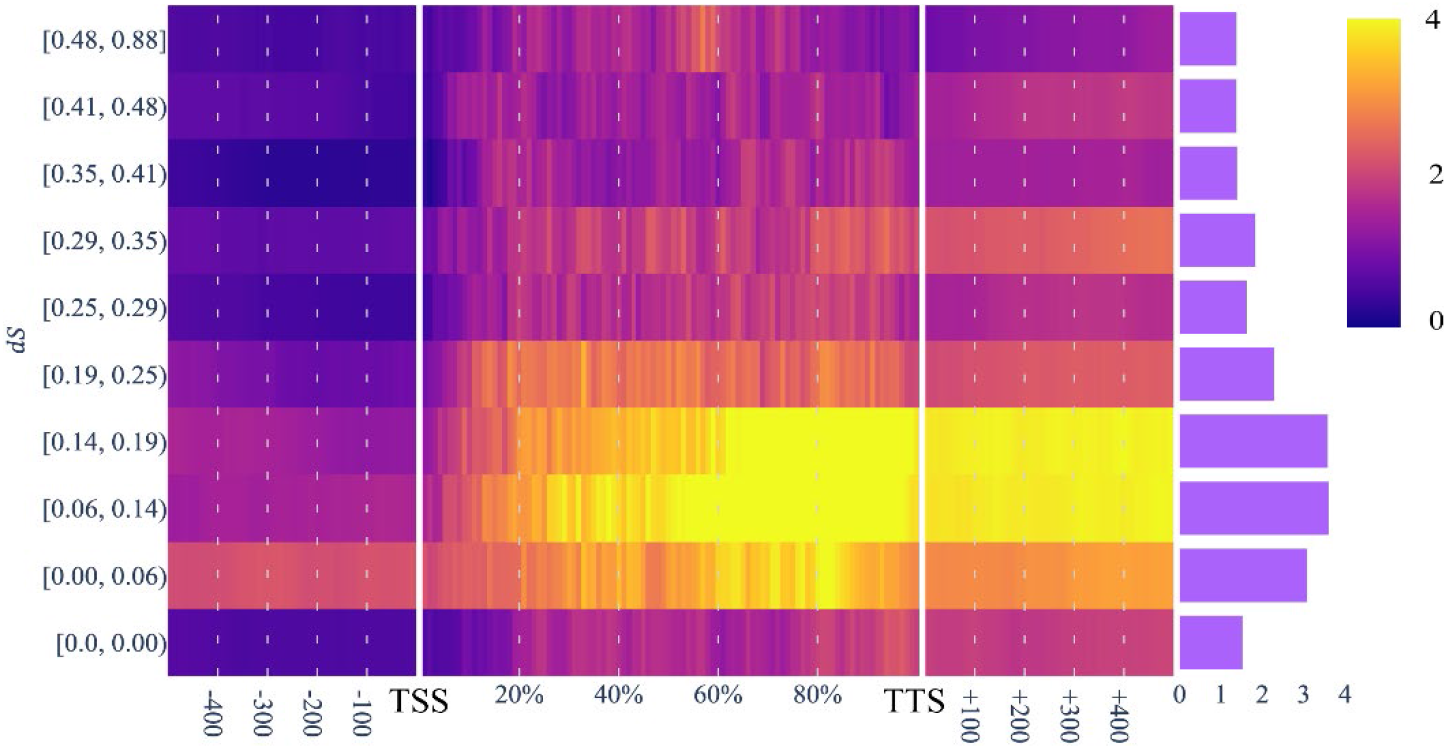
Heatmap of H3K9me3 enrichment in human paralogs broken up into *dS* deciles (y-axis), with the bar graph on the right representing enrichment within the predictive regions.

As genes with an overlapping H3K9me3 peak in *D. discoideum* were inordinately close to TEs, we examined whether the human genes covered by the repressive mark were also close to TEs. We started by examining the proportion of TE sequences that overlapped an H3K9me3 peak in comparison to sequences of endogenous coding genes. Approximately 11% of TE sequences overlapped an H3K9me3 peak, with 1.9% of all TE sequence bases covered by H3K9me3. Interestingly, while ∼42% of coding gene sequences (including introns) from the GRCh38 assembly overlapped an H3K9me3 peak, only 1.1% of all bases within coding gene sequences were covered by H3K9me3, and less than 1% of those bases were in exons. When looking at the set of independent duplicate pairs and singletons separately, we found that 1.5% of exon bases overlapped an H3K9me3 peak in the duplicates, which was significantly higher than singletons in which only 0.24% of exon bases overlapped an H3K9me3 peak (*p* value < 0.001, Mann-Whitney U).

### Triplosensitive human duplicates not enriched for H3K9me3 but enriched for TE sequences

Using dosage-sensitivity predictions published by (Collins et al. 2022), we next sought to test if the repressive mark H3K9me3 was specifically enriched in triplosensitive duplicates, as would be suggested by the dosage balancing model. We found that the probability of triplosensitivity among young duplicates overlapping an H3K9me3 peak was not significantly greater than among those that did not overlap an H3K9me3 peak (supplementary figure 7a, *p* value 0.26, Mann-Whitney U). However, we found that the probability of triplosensitivity was significantly higher among paralogs that overlapped a TE than those that did not (supplementary figure 7b, *p* value < 0.001, Mann-Whitney U). Interestingly, this association was found for three out of the four major TE families present in humans: LINEs, LTRs and DNA (class II) transposons, but was not found with SINEs.

## Discussion

### Duplicate genes show rapid epigenetic de-activation followed by increasing activation over evolutionary time

In the present study, we started by establishing the relationships between key chromatin features and gene expression in *D. discoideum*, expanding on part of the analysis conducted by Wang et al. (2020). Several histone modifications and the presence of open chromatin correlated with gene expression in ways typical of most eukaryotes (Kouzarides 2007; Zhao and Garcia 2015). H3K4me3, H3K27ac, and open chromatin all increased in enrichment within the gene body of coding genes with increasing expression, and H3K9me3 showed enrichment broadly across the gene body and surrounding regions in the lowest expressed genes, with the primary target appearing to be TEs (figure 1a-d, supplementary figures 1, 2). Enrichment for these features followed a pattern that closely tracked the changes in expression as duplicates diverged, suggesting that epigenetic modifications may be a key factor contributing to the rapid reduction of expression in genes soon after duplication, as well as the subsequent expression increase over time of duplicate genes that are preserved. This is consistent with studies in animals that have shown a general trend of higher levels of repressive DNA methylation in young duplicates along with lower levels of overall expression when compared to orthologous singletons (Keller and Yi 2014; Chang and Liao 2012; Huang and Chain 2021). Additionally, our finding of enrichment for the repressive histone modification H3K9me3 in young duplicates is consistent with the enrichment for this mark found in younger duplicates in *C. elegans* and *D. melanogaster* (Chang and Liao 2017), and might be a contributing factor in other lineages such as fish, that appear to lack an association with DNA methylation at the very earliest stages after duplication when paralogs are copy-number polymorphic (Chain et al., 2024).

One key difference with the trend found in these studies, however, was our finding of a mixed population of young duplicates with respect to chromatin modifications and expression; many of the youngest duplicates appear to be enriched for activating modifications, and accordingly show gene expression levels closer to the average among all coding genes, while others appear to be repressed by H3K9me3. For these young duplicates with higher levels of gene expression and activating chromatin modifications, it is possible that this initial expression increase was the result of a transient positive selection pressure related to specific environmental conditions (Huang et al. 2019; Kondrashov 2012; Axelsson et al. 2013). The activating modifications may also merely be present in a way that reflects the ancestral state and has a neutral effect on fitness, with the cis-regulatory sequences important for their recruitment subsequently removed over time via drift. In either case, as duplicate genes age, they seem to experience a rapid decrease in both average expression and coverage by activating modifications before increasing over time (figure 2a-c). One potential explanation for this decrease in expression is the gradual accumulation of cis-regulatory element mutations that reduce the binding affinity of components of the transcriptional machinery. While this explanation fits our results, it is also possible that other epigenetic repressive modifications that we did not measure, like H3K27me3, were recruited to those loci in the early stages of divergence.

### H3K9me3 enrichment is closely associated with TE proximity and repression among younger duplicates

The young duplicate genes with enrichment for the repressive mark H3K9me3 were found to be inordinately close to TEs (figure 3c) which we found to have a strong correlation with the repressive mark in *D. discoideum* (supplementary figure 1). Given the fact that it has been established that H3K9 methylation can spread beyond the target repeat sequence at which it was first nucleated (Choi and Lee 2020; Wang et al. 2014), it seems likely the enrichment for H3K9me3 is the result of the inadvertent spread of this mark to neighboring regions, rather than the deliberate targeting of some young duplicate genes. Importantly, as H3K9me3 enrichment was strongly linked with transcriptional repression in these young duplicates (figure 3c), this could provide a dosage balancing mechanism early in their evolution were they to be born near a TE, or if a TE inserted near a recently duplicated gene. In humans we found as well that younger duplicates were inordinately enriched for H3K9me3, and that this enrichment correlated with TE proximity, suggesting a potentially conserved pattern of early duplicate gene regulation that is shared among species that diverged more than a billion years ago. Surprisingly, while triplosensitivity did not correlate with H3K9me3 enrichment, we nonetheless found that triplosensitive paralogs were inordinately close to TEs. While repetitive sequences can increase the probability of sequence duplication, owing to unequal crossing over between homologous strands during meiotic recombination (Shaw et al. 2002; Jurka 2004), this would not explain the propensity for dosage sensitive duplicates specifically to be enriched near TEs. Instead, it is possible that the spread of another repressive modification associated with TEs in humans, possibly cytosine methylation or H3K9me2, has created a heterochromatic environment favorable for the dosage sensitive duplicates. In the tea plant, for example, it has been found that duplicate genes with intronic TE sequences are enriched for repressive methylation (Tong et al. 2021). It would potentially be illuminating to test for the enrichment of these modifications in triplosensitive duplicates in humans, and to expand our analysis to other organisms to investigate the degree to which TE-mediated repression of young duplicates is conserved across eukaryotic lineages.

## Methods

### Identifying duplicate genes and transposable elements in D. discoideum

To identify duplicate genes in *D. discoideum*, we selected genes classified as paralogs by the Ensembl Compara pipeline (Howe et al. 2021) in release 52 of Ensembl Protists (Howe et al. 2020), excluding TEs, non-coding RNAs, pseudogenes, and gene pairs with a paralogy type annotated as ‘gene split’. To create a set of independent paralog pairs, pairs were filtered to those sharing a reciprocal highest average percent identity. To avoid saturation of synonymous mutations between paralog pairs, only those with a pairwise dS <= 3 were kept. This left us with 953 paralog pairs (supplementary table 1).

We used RepeatModeler2 (Flynn et al. 2020) to predict the presence of transposon sequences in the reference genome. We then used the union of the predicted TEs and those previously annotated in the dictyBase reference (Fey et al. 2013) for all subsequent analyses (supplementary table 2).

### Identifying duplicate genes and transposable elements in humans

We identified human duplicates by selecting genes labeled as paralogs by the Ensembl Compara pipeline in Ensembl release 99 (Howe et al. 2021), excluding non-coding RNAs, pseudogenes, and gene pairs with a paralogy type annotated as ‘gene split’. To create a set of independent paralog pairs, pairs were filtered to those sharing a reciprocal highest average percent identity. To avoid saturation of synonymous mutations between paralog pairs, only those with a pairwise dS <= 3 were kept. This left us with 4029 paralog pairs (supplementary table 3).

Repeat sequence annotations were taken from the UCSC genome browser RepeatMasker track on the GRCh38.p13 assembly. For all analyses, repeat sequences identified by UCSC as transposons but with an uncertain class (i.e., with a class name appended with “?” in the UCSC track data) or a class of “unknown” were excluded. The remaining TEs were categorized as LINEs, SINEs, LTRs, or DNA-transposons.

### ChIP-seq and ATAC-seq alignment and peak calling in D. discoideum

Raw ChIP-seq and ATAC-seq FASTQ reads from a previous experiment of the axenic A2 strain of *D. discoideum* by Wang *et al*. (Wang et al. 2021) (accessions listed in supplementary table 4) were trimmed for adapters and quality using Trim Galore (Kreuger 2023) using default parameters. Filtered reads were aligned to the *D. discoideum* AX4 reference genome from Ensembl Protists release 52 (Eichinger et al. 2005; Howe et al. 2020) using Bowtie2 (version 2.5.2) with default parameters (Langmead and Salzberg 2012). Duplicate reads were removed with Picard MarkDuplicates (version 2.26) (Broad Institute 2019).

An initial set of candidate peaks were called for each ChIP-seq dataset using MACS3 (Zhang et al. 2008) with input DNA samples used as negative controls, and with a liberal *p* value cutoff of 0.10. The irreproducible discovery rate (IDR) (Li et al. 2011) was then calculated for all peaks from each pair of replicates using a Python implementation (Boleu et al. 2021). Replicate peaks with an IDR <= 0.05 were kept for downstream analysis. For ATAC-seq peaks, MACS3 was used with a more liberal *p* value cutoff of 0.3 and the added parameters with parameters ‘--shift -100 --extsize 200’ to set the fragment length to 100 and to smooth the pileup, respectively. Replicates of samples with a Spearman rank correlation less than 0.5 or a low fraction of peaks with an IDR less than 0.05 in any of the three life-cycle stages used (“fruiting body”, “vegetative”, “mound”) were removed from the analysis, along with samples corresponding to the same histone modification in the remaining life-cycle stages. All samples from the “streaming” stage were removed, as both H3K27ac and H3K9me3 replicates had a low number of peaks with IDR less than 0.05 or a low Spearman correlation, respectively.

### ChIP-seq and ATAC-seq enrichment in D. discoideum

To measure the association between gene expression and enrichment for each ChIP-seq and ATAC-seq peak at each point along a typical gene, we first binned all protein coding genes (with the exception of TEs) into deciles based on expression averaged across all samples from each life cycle stage. We next calculated the frequency of each peak type at each base from 500 bases upstream of the TSS to 500 bases downstream of the TTS using all genes within each expression decile. For each gene, the peak data overlapping the gene body was divided into percentiles to create a common basis of comparison despite varying gene body lengths. The per-base frequencies were then divided by the average frequency across all bases in all protein coding genes.

When measuring the association between enrichment for ChIP-seq and ATAC-seq peaks and duplicate pair divergence, we separated all independent duplicate pairs with a *dS* <= 3 into *dS* deciles. Again, we calculated the frequency of each peak type from TSS-500 to TTS+500 for all duplicates within each decile. The per-base frequencies were then divided by the average frequency across all bases in singletons to provide a fold-change over the singleton average.

### H3K9me3 enrichment in humans

A total of 16 primary cell-based samples from four tissues in four individuals were used for the human H3K9me3 data. Data consisted of processed H3K9me3 ChIP-seq peaks accessed via ENCODE [cite]. For details on the pipeline used by ENCODE when processing the H3K9me3 ChIP-seq data, see the pipeline protocol published alongside the raw and processed for each accession listed in supplementary table 5. The processed peak calls were used to calculate a fold-change in enrichment among human paralogs over the average for singletons in the same way as done for the *D. discoideum* data.

### Mapping dosage sensitivity predictions to GRCh38

A table of triplosensitivity and haploinsufficiency probabilities was previously generated by the authors of (Collins et al. 2022). Genes included in that data set consisted of canonical transcripts from Gencode v19, corresponding to the human genome assembly GRCh37. Given the H3K9me3 peak data, as well as *dS* estimates were based on sequences from GRCh38, only genes that could be lifted over from GRCh37 with at least 50% of their sequence contiguously mapping to GRCh38 were kept. Lift over of the genome was performed using the ‘rtracklayer’ (version 1.64) (Lawrence et al. 2009) package for the R programming language (version 4.4) (Ihaka and Gentleman 1996).

### RNA-seq alignment and transcript quantification

Raw RNA-seq FASTQ reads from Wang et al. (Wang et al. 2021) were trimmed for quality and adapter removal us Trim Galore with default parameters (version 0.6.1) (Kreuger 2023) and pseudo-aligned to all *D. discoideum* transcripts using Kallisto (version 0.46) (Bray et al. 2016). The reference transcript sequences were derived from Ensembl Protists release 52 (Howe et al. 2020). Expression values were normalized to transcripts per kilobase million (TPM) from the transcript abundances using the Sleuth package (version 0.30) (Pimentel et al. 2017) for the R programming language (version 4.4) (Ihaka and Gentleman 1996).

To adjust gene expression for possible inflation due to the presence of cryptic gene copies not annotated in the reference genome, RNA transcript counts mapping to duplicate loci were adjusted based on the level of ChIP-seq input DNA signal at each gene’s respective locus with the following equation:

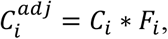

where the adjusted transcript count for the i^th^ paralog 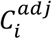 is the product of the original normalized TPM of the i^th^ paralog *C*_*i*_and its associated scaling factor *F*_*i*_, which is defined as the ratio of the average input DNA read pileup per base along the paralog sequence over the averaged input DNA read per base across all single copy genes if *F*_*i*_>1, and 1 otherwise.

## Data and Code Availability

*Dictyostelium discoideum* AX4 reference DNA sequences, genomic feature annotations and reference gene transcript sequences were accessed from Ensembl Protists release 52 (Howe et al. 2020), and annotation of transposable element open reading frames were downloaded from dictybase.org (Fey et al. 2013).

All *D. discoideum* ChIP-seq, ATAC-seq, and RNA-seq data were downloaded as raw FASTQ files from NCBI with BioProject accession PRJNA565988, submitted originally by Wang *et al* (Wang et al. 2021). *D. discoideum* ChIP-seq samples used in this study consisted of two biological replicates for each histone modification and input DNA negative controls from each of three D. discoideum life cycle stages: vegetative, mound, and fruiting body. ATAC-seq and RNA-seq samples also included two biological replicates from each of the above-mentioned stages. For more detailed information about each individual *D. discoideum* sample, including associated GEO accessions, see supplementary table x. All 16 human H3K9me3 peak data samples were accessed via ENCODE, with specific sample accessions listed in supplementary table 5.

Scripts used to process and analyze the data can be found at github.com/tdw-89/wolfe_chain_2024.

## Acknowledgements

We would like to thank and acknowledge Dr. Eric Greer for correspondence related to the data used in this study originally generated by Wang et al. (2021).

